# Activation thresholds set the speed of biochemical trigger waves

**DOI:** 10.64898/2026.10.01.756064

**Authors:** Zhengda Li, James E. Ferrell

## Abstract

Trigger waves are positive feedback-mediated traveling waves that propagate without slowing down or diminishing in amplitude, enabling biochemical coordination over distances where diffusion alone would be insufficient. Apoptosis can spread as trigger waves of caspase activity, as demonstrated in *Xenopus* eggs and egg extracts. We measured apoptotic wave speeds in extracts at two cell cycle states and found that waves propagated more slowly in M-phase arrested cytostatic factor (CSF) extracts than in interphase extracts (21 versus 27 µm min^−1^). Luther’s equation, 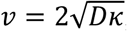, predicts wave speed from the diffusion coefficient *D* and the positive-feedback doubling rate *κ*. However, *D* and *κ* were comparable between the two extracts and could not explain the speed difference. This discrepancy highlighted a key limitation of the classical equation: it does not account for an activation threshold in trigger wave initiation, and in fact the threshold required to trigger apoptosis was found to differ between cell cycle states. We therefore extended Luther’s equation to include such a threshold, obtaining 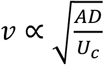, where *A* is the reaction rate and *U* is the activation threshold. Perturbing apoptotic sensitivity confirmed the predicted threshold–speed relationship. These findings establish the activation threshold as a key determinant of trigger wave speed and provide a broadly applicable scaling law for predicting the spread of switch-like biochemical signals.

## Main text

Trigger waves allow biochemical signals to spread through cells and tissues without slowing down or diminishing in amplitude^1–3^. They arise when diffusion, which normally supports only the local spread of an activated signaling molecule, is coupled to positive feedback reactions that generate new active signaling molecules near the wavefront. Familiar examples of biological trigger waves include action potentials^4^ and calcium waves^5^. Apoptotic caspase activation^6^ and mitotic cyclin B-Cdk1 activation^7^ also spread as trigger waves. In all of these cases the trigger waves are confined to single cells, but there are also a number of examples of trigger waves that spread from cell to cell in a tissue. For example, trigger waves of ERK activation spread between cells after wounding in mammalian epithelia^8,9^ and fish scales^10,11^. Bisected planaria also relay their regeneration responses via intercellular ERK trigger waves^12^. And ferroptotic cell death spreads intercellularly via reactive oxygen species-dependent trigger waves^13^. Whether within individual large cells or between cells in a tissue or organism, trigger waves allow all-or-none signals to be relayed rapidly and reliably over distances where diffusion alone would be slow.

The speed of a trigger wave is at least partially determined by two factors: the speed of the spatial spread of the active trigger wave mediators, often quantified by a diffusion coefficient (typically in µm^2^ s^−1^), and the speed of the positive feedback reactions, one measure of which is the doubling time (in s) or apparent first-order rate constant (in s^−1^) for the exponential growth phase of the reactions. Dimensional analysis argues that the wave speed (in µm s^−1^) should then be given by 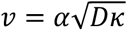 or 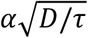, where *α* is a dimensionless coefficient, *D* is the diffusion coefficient, *κ* is the apparent first-order rate constant for the positive feedback, and *τ* is ln2/*κ*. This relationship was first proposed by Robert Luther in 1906^14,15^, and he asserted (without providing justification) that the value of *α* was between 2 and 10 for chemical waves. Further theory showed that if the reaction system has a negligible threshold for activation, the value of *α* will be 2^16–18^. Luther’s equation has proven to be a valuable guide for estimating and understanding the speed of a given trigger wave^7,19,20^.

We have been studying apoptotic trigger waves in *Xenopus* eggs and extracts. Previous work established that apoptosis accounts for the spontaneous ultimate death of unfertilized *Xenopus* eggs, and that it spreads in a rapid, stereotyped fashion from the vegetal pole to the animal pole of the egg^6^. This process can be recapitulated in cell-free interphase *Xenopus* egg extracts: if one part of an extract is made apoptotic by treatment with cytochrome c, caspase 3/7 activation and apoptosis propagates into and through the rest of the extract at a constant speed of about 30 µm min^−1^ or 0.5 µm s^−1 6,20^. This allows apoptosis to rapidly spread throughout a cell, provided that the cell has a simple geometry, as the spherical *Xenopus* egg does. However, more complicated geometries can generate other behaviors. For example, if apoptotic waves are initiated in a cytoplasmic tube of less than a few microns’ diameter, they will be arrested at the junction of the tube with a large cytoplasmic chamber^21^. The arrest of the wave occurs because the diffusing apoptotic mediators are diluted more effectively at the junction than they were in the thin tube, and this dilution extinguishes the propagating wave. The fact that trigger waves can be geometrically compartmentalized may help explain why caspase activation remains localized to individual dendritic spines when a neuron undergoes synaptic pruning^22^.

Our previous work on apoptosis in *Xenopus* extracts mainly employed interphase-arrested extracts, in part because it had been reported that the mitotic protein kinase cyclin B-Cdk1 can negatively regulate the apoptotic control system^23–25^. Here we have examined apoptosis in both interphase extracts and cytostatic factor-(CSF-) arrested extracts, which have higher levels of cyclin B-Cdk1 activity and which mimic the cell cycle state of the unfertilized egg. We found that apoptosis could still propagate via trigger waves in CSF-arrested extracts, and that the speed was about 20% lower than in interphase extracts. However, this slowing was not accounted for by Luther’s equation; both the exponential doubling speed for caspase 3/7 activation and the diffusion coefficient for a protein-sized probe were the same in CSF-arrested extracts and interphase extracts. This led us to reconsider the assumptions that underpin Luther’s equation. Here we analytically derive an extended version of Luther’s equation that takes into account the possibility of a non-negligible threshold in the reactions underlying a trigger wave, using a piecewise-linear model of the reaction rate equation. By numerical simulation we then show that this equation holds for a more realistic model that cannot be solved analytically. Finally, through experiments with drugs, peptides, and proteins that make extracts more or less sensitive to apoptotic stimuli, we show that this theoretical relationship between threshold and wave speed holds for apoptotic trigger waves in *Xenopus* egg extracts.

## Results

Previous studies have suggested that the master regulator of mitotic entry, cyclin B-Cdk1, may be a key modulator of apoptosis. Active Cdk1 has been reported to inhibit apoptosis through the phosphorylation of caspase-2^24^, caspase-9^23^ and BIRC5^25^. However, elevated Cdk1 activity has also been reported to promote apoptosis through the phosphorylation of BAD^26^, MCL1^27^ and XIAP^28^. Here we set out to determine if the dynamics of apoptosis differs in *Xenopus* egg extracts arrested in meiosis versus interphase.

We prepared undiluted CSF-arrested and interphase-arrested extracts by standard methods (see Methods)^29,30^ except that we included nocodazole in both types of extracts to make the extracts more spatially homogeneous^6,31,32^ (Fig. 1a). To monitor caspase activation we added Z-DEVD-R110, a fluorogenic substrate for caspase-3/7, and we also included the proteasome inhibitor bortezomib to decrease the background levels of proteasome-mediated cleavage of the caspase probe^6,21^. The extracts were loaded into Teflon tubes, and apoptosis was initiated by dipping one end of the tube into an extract that had been made apoptotic by treatment with cytochrome c, as previously described^6^ (Fig. 1b). The tube was placed under mineral oil and the spatial propagation of caspase activity was monitored by fluorescence imaging, again as previously described^6^.

**Fig. 1:**
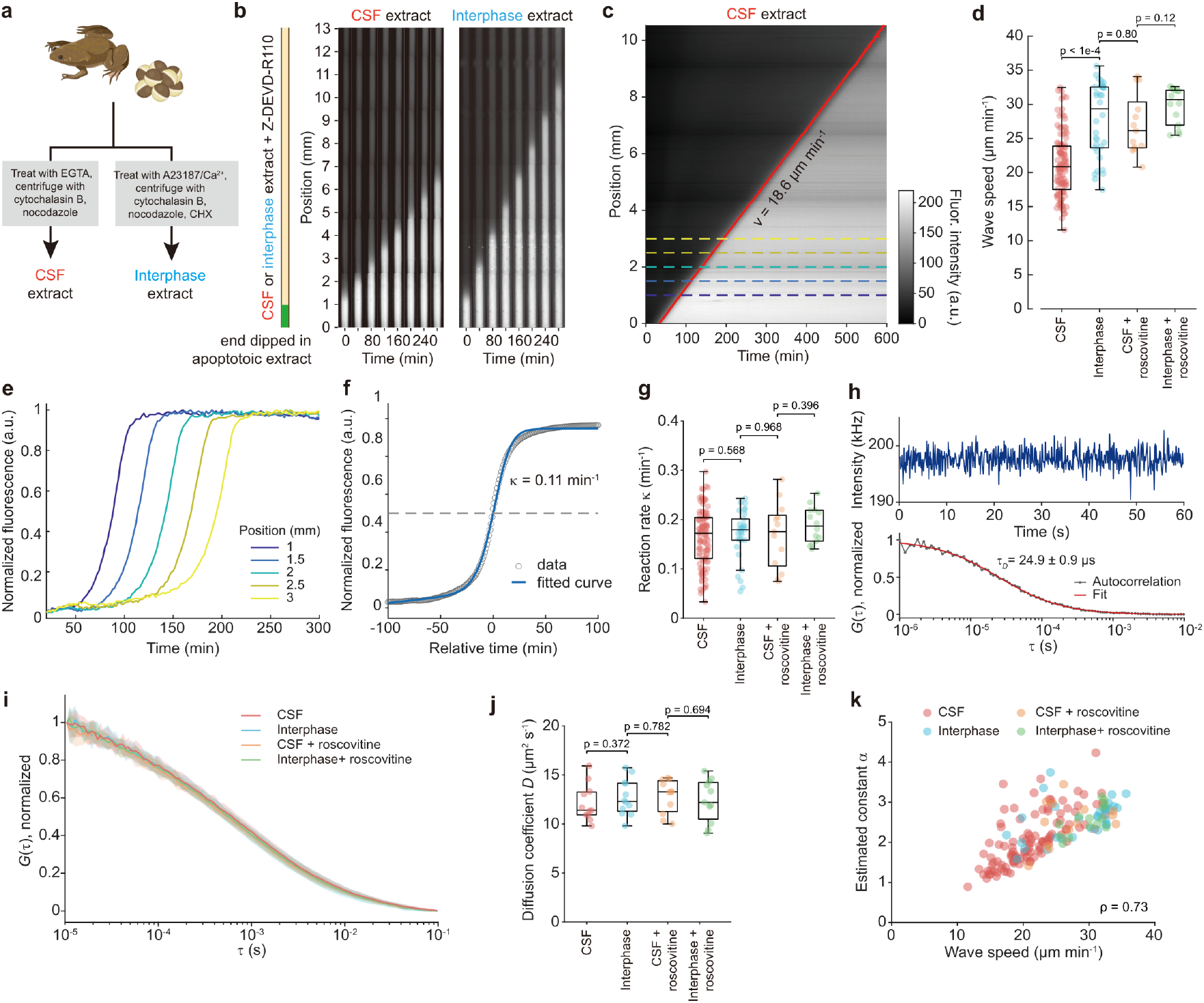
Apoptotic waves propagate at different speeds in interphase and CSF-arrested *Xenopus* egg extracts. a, Schematic illustration of the preparation of *Xenopus* egg extracts arrested in different cell cycle stages. b, Apoptotic trigger waves. CSF and interphase *Xenopus* egg extracts containing the fluorogenic caspase-3/7 sensor Z-DEVD-R110 were loaded into Teflon tubes and one end of the tube was dipped into activated apoptotic extract to initiate apoptosis. The resulting apoptotic wave was followed by fluorescence video microscopy. Snapshots of the tubes at 40 min intervals were assembled into the panels shown here. c, Representative kymograph showing an apoptotic wave in a CSF-arrested extract. The linearity of the wavefront indicates a constant propagation speed. The red line indicates the contour corresponding to the half maximal fluorescence intensity at each position. A linear fit of this contour yielded a speed of 18.6 μm min^−1^. The dashed lines indicate the positions at which the time courses in panel e were taken. d, Apoptotic wave speeds in CSF extracts (red, N = 113 replicates from 43 frogs), interphase extracts (blue, N = 34 replicates from 12 frogs), CSF extracts treated with 10 μM roscovitine (orange, N = 13 replicates from 4 frogs), and interphase extracts treated with 10 μM roscovitine (green, N = 14 replicates from 5 frogs). P-values from the rank-sum test are as indicated. e, Temporal dynamics of normalized fluorescence intensity in an CSF-arrested extract at five positions in the tube. The time courses shown here were obtained at positions shown in Fig. 1c. f. Superimposition of several time courses by time-shifting so that the normalized average fluorescence has half-maximal at time zero. Time course data from a CSF-arrested extract sampled at 200 positions were aligned by time-shifting (dots) and averaged, a logistic function was fitted to the data to estimate 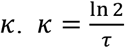 where *τ* is the half-time of exponential growth phase. g, The apoptosis autocatalytic rate constant *κ* from multiple experiments for CSF extracts (red, N = 110 replicates from 40 frogs), interphase extract (blue, N = 30 replicates from 11 frogs), CSF extract treated with 10 μM roscovitine (orange, N = 13 replicates from 4 frogs), and interphase extract treated with 10 μM roscovitine (green, N = 14 replicates from 5 frogs). P-value from rank-sum test is labeled on top. h, A representative fluorescence-correlation spectroscopy (FCS) intensity trace (upper) and normalized autocorrelation curve (lower) used to estimate diffusion time of the diffusion standard ATTO 488.The estimated diffusion time is 24.9 ± 0.9 μs (mean ± std). i, Overlay of average normalized FCS autocorrelation curves for Alexa-488 conjugated bovine serum albumin from CSF (red, N = 11 replicates from 5 frogs), interphase (blue, N = 12 replicates from 5 frogs), CSF + roscovitine (orange, N = 11 replicates from 5 frogs) and interphase + roscovitine extract (green, N = 11 replicates from 5 frogs). Shaded regions represent standard deviation among replicates, showing all four conditions have similar autocorrelation profiles. j, Effective diffusion coefficients for Alexa 488-conjugated bovine serum albumin in CSF extracts (red, N = 11 replicates from 5 frogs), interphase extracts (blue, N = 12 replicates from 5 frogs), CSF extracts treated with roscovitine (orange, N = 11 replicates from 5 frogs), and interphase extracts treated with roscovitine (green, N = 11 replicates from 5 frogs). P-values from the rank-sum test are shown. k, Relationship between the measured apoptotic wave speed and the Luther’s equation scaling constant *a* for different types of extracts (color-coded). A positive correlation (Spearman’s ρ = 0.73) was observed.

Snapshots from one such experiment are shown in Fig. 1b. Apoptosis was detected in both the CSF-arrested extract and interphase extract, and in both cases the fluorescent wavefront proceeded toward the far end of the tube at a constant speed over distances of several millimeters and times of several hours (Fig. 1b). The imaging data were made into kymographs (e.g. Fig. 1c), which were fitted to estimate the wave speed. The apoptotic wave speeds were 20-25% slower in CSF-arrested extracts than in interphase extracts (Fig. 1b-d); average speeds from multiple independent extract preparations were 21 ± 5 μm min^−1^ (mean ± standard deviation) for CSF extracts versus 27 ± 5 μm min^−1^ for interphase extracts (Fig. 1d). The lower speed seen in CSF extracts was rescued by addition of 10 μM roscovitine, a Cdk inhibitor, whereas roscovitine had no apparent effect on the speed of apoptotic waves in interphase extracts (Fig. 1d). Taken together, these data show that apoptotic trigger waves can propagate in both types of extract and are slowed by Cdk activation.

It seemed plausible that the slower wave speed in CSF extracts was due to a decrease in the doubling speed for the positive feedback reactions that regulate the production of active caspase 3/7. From Luther’s equation, a ~40% reduction in the doubling speed (*κ*) would account for the observed ~22% decrease in the wave speed. To test this hypothesis, we measured the apparent doubling speed of the apoptotic reactions using an approach described previously^20^. We normalized the time courses of Z-DEVD-R110 cleavage at each position along the tube (Fig. 1c,e), shifted each trace so that its half-maximum time aligned across positions, and fitted the averaged activation profile with a model in which caspase-3/7 activity grows logistically (Fig. 1f, see methods). The fitted exponential growth rate gives the doubling speed *κ*. Contrary to our expectation, *κ* was not significantly lower in CSF extracts than in interphase extracts, and 10 μM roscovitine had no detectable effect on *κ* in either extract type (Fig. 1g). Thus, the slower wave speed in CSF extract cannot be explained by slower exponential amplification of caspase activity.

We next asked whether the cell-cycle dependence of wave speed could reflect differences in cytoplasmic diffusion. Using fluorescence correlation spectroscopy (FCS) with Alexa Fluor 488-conjugated bovine serum albumin (BSA), and ATTO 488-carboxylic acid as a calibration standard (*D* = 400 μm^2^ s^−1^, Fig. 1h), we estimated protein diffusivity in each extract condition (Fig. 1i; Methods). The fitted diffusion coefficients were similar in CSF and interphase extracts (12.1 ± 1.9 µm^2^/s and 12.7 ± 1.8 µm^2^/s, mean ± SD), and roscovitine did not measurably alter the diffusivity (Fig. 1i,j). Therefore, neither *κ* nor *D* accounts for the slower apoptotic wave speed in CSF extract.

We then calculated the dimensionless pre-factor 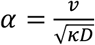 for each experiment, which is commonly assumed to equal 2^14,17^. The mean value of *α* was found to be close to 2 (2.23 ± 0.64, mean ± SD), consistent with previous analyses of mitotic and apoptotic trigger waves^7,17,20^, but estimates from individual experiments varied from approximately 1 to 4. The estimated values of *α* correlated strongly with the measured wave speed (Spearman’s ρ = 0.73, p = 3.3e-28, *α* = 2.1 ± 0.65 for CSF and *α* = 2.6 ± 0.49 for interphase extract, Fig. 1k). Treating *α* as a constant is therefore too restrictive for these data; additional variables not considered in Luther’s equation appear to modulate apoptotic wave speed across cell cycle states.

### CSF and interphase extracts differ in the threshold concentration of cytochrome c required for apoptosis

A plausible missing parameter is the activation threshold of the apoptotic switch. Apoptosis requires upstream input, such as the release of cytochrome c from the mitochondria, for activation^6^, and previous work has shown that the cytochrome c concentration must exceed a critical level to produce detectable amounts of caspase activation^33–35^. Such a threshold suggests that the apoptotic control system is a bistable switch, with the threshold corresponding to the concentration of caspase where the system traverses a saddle-node bifurcation and flips from a stable low-caspase activity steady state to a stable high-caspase activity state. This would mean that as a wavefront of caspase activity progresses through space, three factors would contribute to the wave speed: the speed of diffusion; the value of the threshold that the active caspase concentration must exceed before the positive feedback reactions become effective; and finally the speed of the positive feedback, which is relevant once the threshold is exceeded. However, Luther’s equation takes into account only two of these three factors. As pointed out by Showalter and Tyson, both the form of Luther’s equation and the constant value of *α* = 2 follow if it assumed that any threshold is negligible. This assumption allows Luther’s equation to be derived by the approach taken by Fisher, Kolmogorov, Petrovsky, and Piskunov in their analysis of traveling waves in simple unstable (rather than bistable) reaction-diffusion systems^16–18^, and this is why the form and the pre-factor of *α* = 2 in Luther’s equation and the Fisher-KPP equation are the same. We therefore tested whether *Xenopus* extracts become apoptotic only above a critical cytochrome c concentration, whether this threshold differs between CSF and interphase extracts, and whether changes in the threshold might explain the observed wave speed variation.

To measure the cytochrome c vs. caspase dose-response relationship, we added a range of concentrations of cytochrome c to extracts and monitored Z-DEVD-R110 cleavage. In a representative CSF extract, caspase was maximally activated by 20 min for cytochrome c concentrations of 80 nM or more but was never measurably activated for 70 nM or less, implying a critical concentration *U*_*c*_ of between 70 and 80 nM (Fig. 2a). These concentrations varied from extract to extract, but overall the interphase extracts were activated at lower cytochrome c concentrations than CSF extracts (Fig. 2b). By logistic fitting to all of the extract data, the threshold *U*_*c*_ was 55 ± 4 nM in interphase extracts and 66 ± 3 nM in CSF extracts (Fig. 2c). The total cytochrome c concentration in *Xenopus* eggs has been estimated to be 1 µM^36,37^, so these threshold concentrations represent 5-7% of the total.

**Fig. 2:**
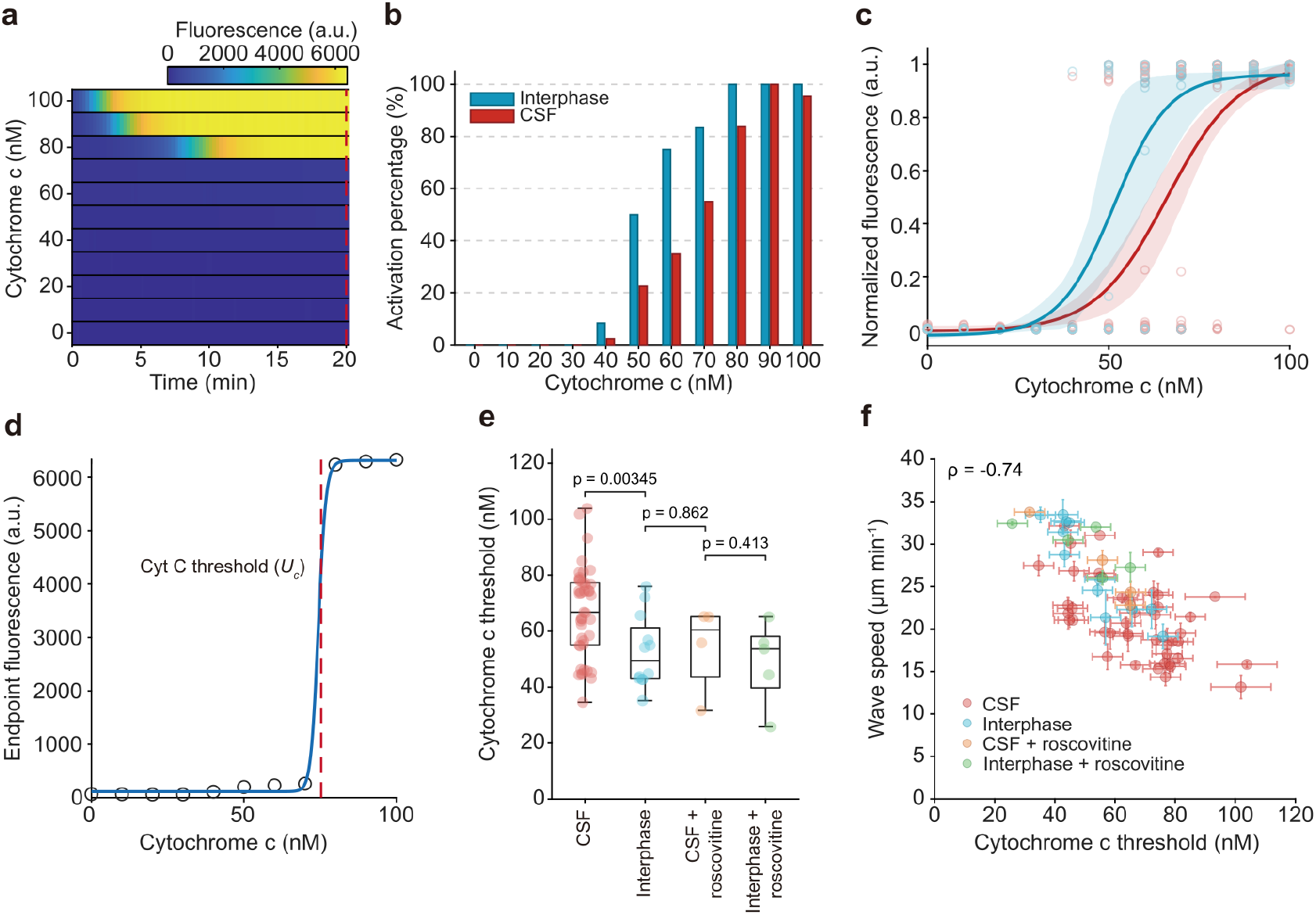
Cytochrome c thresholds in interphase versus CSF *Xenopus* egg extracts. a, Temporal dynamics of sensor cleavage after treating a CSF *Xenopus* egg extract with several concentrations of cytochrome c. The fluorescence intensity as a function of time is shown as a color-coded heat map. b, Percentage of extracts exhibiting apoptosis as a function of cytochrome c concentration in CSF-arrested (red) and interphase (blue) extracts. Data were collected from 43 frogs. For each frog, the egg extract was divided into aliquots and supplemented with increasing concentrations of cytochrome c. Caspase sensor fluorescence was measured after 20 min, and apoptotic activation was defined as fluorescence exceeding 50% of the maximum value observed within that experiment. c, Logistic fits of caspase sensor data at 20 min as a function of cytochrome c concentration for CSF (red) and interphase extracts (blue). The shaded regions represent the 95% confidence interval of the fitted curves. Data is from 43 frogs for CSF extracts and 12 frogs for interphase extracts. d, Estimating the experimental threshold concentration *U*_*c*_ by fitting a Hill function to the dose-response data. R110 fluorescence was measured 20 min after addition of cytochrome c. e, Caspase activation thresholds in CSF extracts (red, N = 43 frogs), interphase extracts (blue, N = 12 frogs), CSF extracts treated with 10 μM roscovitine (orange, N = 4 frogs), and interphase extracts treated with 10 μM roscovitine (green, N = 5 frogs). P-values are from the rank-sum test. f, The relationship between apoptotic wave speed *v* and cytochrome c activation threshold *U*_*c*_. The error bars for the wave speeds represent the standard deviations from 1 to 4 technical replicates, and the error bar is 0 for frogs with only one measurement. The error bars of the cytochrome c threshold represent the step size of the cytochrome c concentration in the experiment. The experimental conditions are color coded as indicated.

We also quantified *U*_*c*_ for individual extracts (Fig. 2d), which confirmed the difference in threshold between CSF extracts and interphase extracts (Fig. 2e). Roscovitine lowered the *U*_*c*_ value in CSF extracts to approximately the interphase level, but had little effect on interphase extracts (Fig. 2e). Finally, *U*_*c*_ was strongly negatively correlated with wave speed (Spearman ρ = −0.74, p = 2.7e-12, Fig. 2f), suggesting that the cytochrome c activation threshold is a strong candidate control parameter for apoptotic trigger wave speed.

We also examined the relationship between the rate of caspase-3/7 activation throughout an apoptosis time course and the inferred concentration of activated caspase present at that time. As shown in Fig. 3a, after cytochrome c treatment, the fluorescence generated by the caspase reporter increased sigmoidally (Fig. 3a). For further analysis, we focused on the central, rapidly changing portion of each trace, where fluorescence was between 20% and 80% of its maximum (Fig. 3a). This minimized contributions from nonspecific cleavage, diffusion, and substrate depletion. Within this range we estimated the normalized caspase-3/7 reaction rate from the substrate cleavage rate (inferred from the fluorescence as described in the Methods section) divided by the remaining substrate concentration. The rate of caspase activation was found to initially increase with time and then plateau (Fig. 3b; Methods). A phase plot of the normalized caspase-3/7 activity 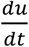 versus *u* showed a greater than linear dependence at low *u* values, a maximum at approximately *u* = 0.3 and a decrease at high *u* values (Fig. 3c, gray circles and blue curve). These findings suggest that positive effects of *u* on 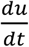 are due to an ultrasensitive Hill-type activation process^21^. The negative effects at high *u* values could be due to depletion of inactive procaspase-3/7 (Fig. 3d).

**Fig. 3:**
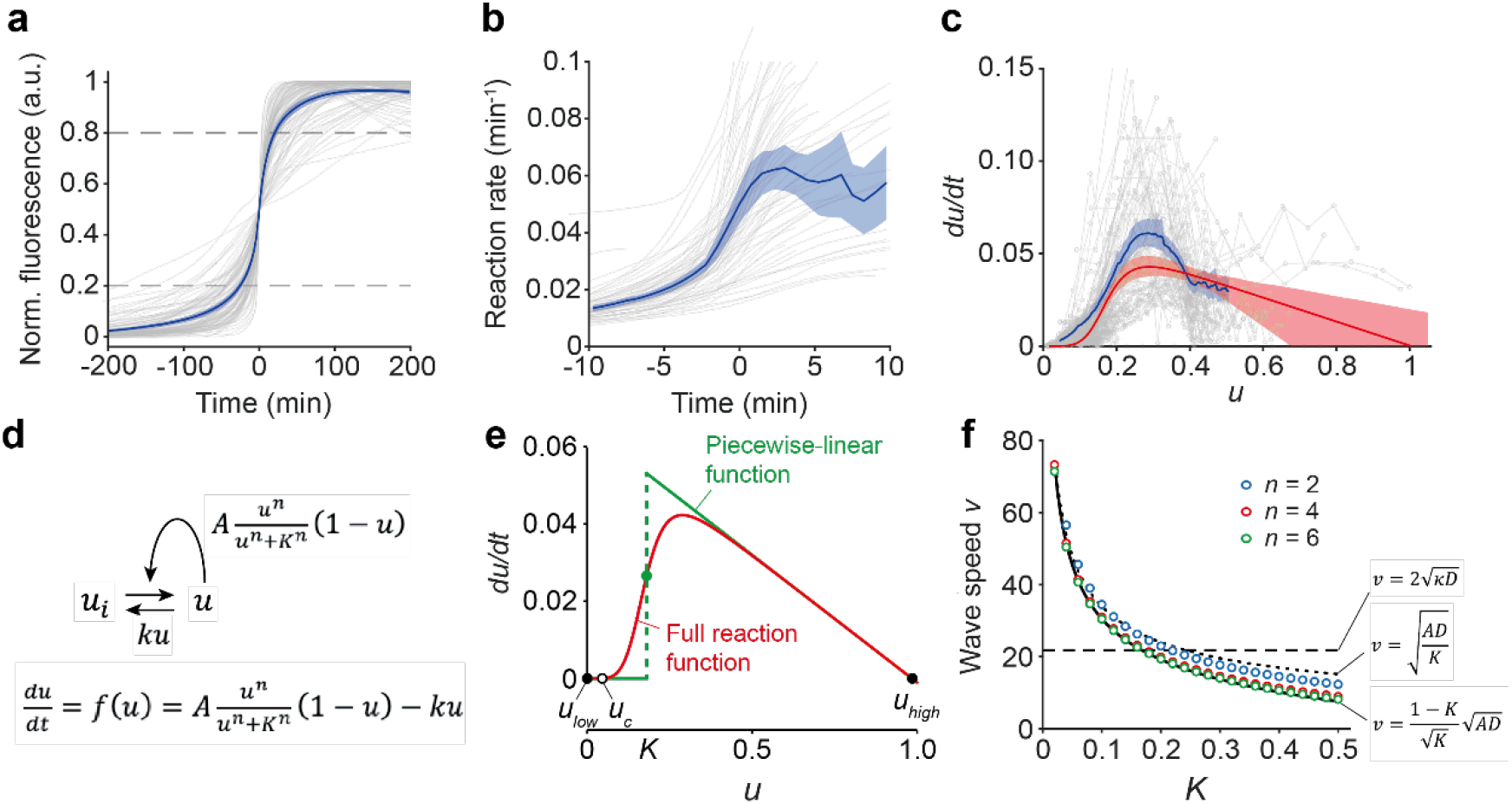
Estimate apoptotic wave speed as a function of the activation threshold. a, Fluorescence dynamics of Z-DEVD-R110 for all experiments with CSF extracts (gray lines, N = 117). The traces have been time-shifted to make the fluorescence be half-maximal at *t* = 0. The blue curve and the shaded area show the average time course and 95% confidence interval respectively. Data between the 0.2 and 0.8 cutoffs (dashed lines) were used for further analysis (N = 117 replicates from 43 frogs). b, Caspase-3/7 first order reaction rate constants, for CSF extracts induced to undergo apoptosis, calculated from the slopes of the time course curves shown in panel **a** divided by the available substrate, taken as (1 – fluorescence intensity). Only the portions of the time courses between the 0.2 and 0.8 cutoffs were analyzed. The gray curves indicate reaction rates as a function of time for each experiment, N = 117. The blue curve and shaded area show the average reaction rate as a function of time and the 95% confidence interval (N = 117 replicates from 43 frogs). c, Phase plot of caspase-3/7 activity (*u, x*-axis) versus its derivative (*du*/*dt, y*-axis) for each individual experiment with CSF extracts (gray circles, N = 1811 points from 117 traces from 43 frogs). The blue curve and the shaded area show the average phase plot and 95% confidence interval. The red line and shaded area represent the fitted curve for the parameters *n* = 5.2 (95% CI 4.4 to 6.2), *K* = 0.18 (95% CI 0.15 to 0.21), *k* = 1.1*e* − 5 (95% CI 1*e* − 6 to 3.0*e* − 3) and *A* = 0.065 (95% CI 0.050 to 0.079), along with the 95% confidence interval around the fitted curve. The caspase activity (*u*) is normalized such that the maximum equals 1. d, A one-variable ODE model for the activation of an apoptotic mediator like caspase-3/7. Note that for simplicity it is assumed that active caspase-3/7 can be inactivated back to inactive procaspase-3/7. In actuality, active caspase-3/7 is removed by degradation and procaspase-3/7 is replaced by new synthesis. Nevertheless this simplification suffices for the purposes of the present analysis, and allows the system to be reduced to a single ODE. e, Approximation of the reaction rate function with Hill exponents (red curve) or with a piecewise-linear function (green curve). The assumed parameters are *K* = 0.18, *n* = 5.2, *A* = 0.065, and *k* = 0.001. For the rate function with Hill exponents, 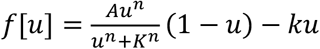. For the piecewise-linear function, 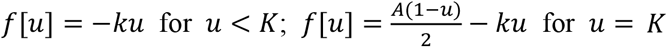; and *f*(*u*) = *A*(1 − *u*) − *ku* for *u* > *K*. The two stable steady states (SSS) for the system are represented by solid circles and the unstable steady state (USS) by an open circle. f, Wave speeds numerically calculated from Equation 2 for systems with varying Hill coefficients (*n*) and activation thresholds (*K*) (circles), compared with the theoretical relationship derived for the piecewise-linear reaction rate function (solid black curve); the simplified theoretical relationship for small values of *K* (dotted black curve); or Luther’s equation (dashed black curve). The other assumed parameters were *A* = 0.17 (from Fig. S2d), *D* = 11 (from Fig. 1j), *k* = 1.7 × 10^−4^.

### Wave speed in a simple model system with a threshold

With these data in hand, we constructed a simple one-variable reaction-diffusion model for a bistable reaction system with an appreciable threshold, with the aim of deriving how the trigger wave speed varies with the position of the threshold. We started with a reaction function that describes the dynamics of a well-mixed positive feedback reaction system wherein an inactive species *u*_*i*_ can be autocatalytically activated by active *u*, and can be inactivated by a mass action process (Fig. 3d):

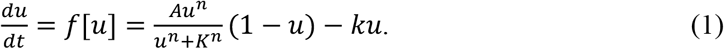

Here *u* is a normalized (ranging from zero to 1) general apoptosis-state variable that represents the collective behavior of the apoptotic pathway or the behavior of some rate limiting species. Note that experimentally caspase-3/7 activity was the principal readout of *u*, whereas cytochrome c serves as the experimentally controlled upstream input to drive the system across its activation threshold, so this one-variable model is simpler than the experimental set up. The parameter *A* is the autoactivation rate constant, *K* determines the activation threshold for the positive feedback, *n* is the effective Hill exponent, and *k* is a small inactivation rate constant.

This function captures the basic features of the ^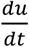^ versus *u* data reasonably well, as shown in Fig. 3c (red curve). The full reaction-diffusion equation for the system is therefore:

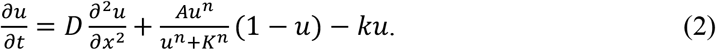

Closed-form traveling wave solutions are difficult to obtain for Hill-type reaction-diffusion systems; further simplification is helpful.^1,38^ Because the fitted Hill coefficient was large (Fig. 3c, red curve), we approximated the Hill activation term by a step function (a Heaviside function) centered on *u* = *K* (Fig. 3e, Equation 3):

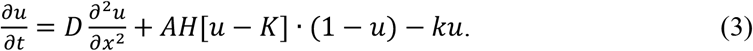

The resulting piecewise-linear approximation to *f*[*u*] (Fig. 3e, green dashed curve) gives an analytical traveling wave solution (Methods). The wave speed depends strongly on *K* and only weakly on *k* if *k* is small (Fig. S1). If we nondimensionalize Equation 3 (Methods), the wave speed is well approximated by:

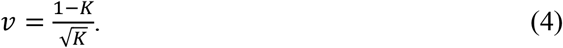

In full dimensional terms, this expression becomes:

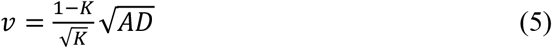

Note that the rate constant *A* plays a similar role to the doubling rate constant *κ* in Luther’s equation. If *K* is much smaller than 1, then the simpler expression:

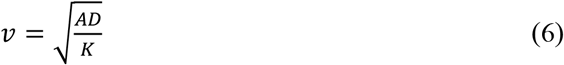

holds. These two equations, plus Luther’s equation (which includes no *K* dependence), are plotted in Fig. 3f.

We compared the two theoretical expressions to wave speeds obtained by numerical solution of the full Hill reaction-diffusion equation for varying values of *K* (Equation 2). The theory agreed well with the numerical results for Hill exponents of 4 or more, and agreed perhaps surprisingly well even for *n* = 2 (Fig. 3f).

### The relationship between threshold and wave speed for apoptotic waves in *Xenopus* egg extracts

To experimentally test the applicability of Equations 5 and 6, we first examined the relationship among three threshold-related quantities: the parameter *K*, which in the Hill function model (Equations 1 and 2) represents the EC50 for caspase-3/7 activation; the unstable steady state *u*_*c*_ for the bistable reaction system (Fig. 3e), which represents the threshold or separatrix of the model (Fig. 3d, e); and experimentally measured *U*_*c*_ which correspond to the cytochrome c activation threshold like those shown in Fig. 2. The quantities *K* and *u*_*c*_ are not equal (Fig. 3e) but they do scale similarly; as pointed out by Schlomann et al., they are related by the expression 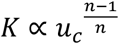 (see Ref. ^39^ and Methods). In the experiments, cytochrome c is the controlled upstream input used to drive the generalized apoptotic state. A cytochrome c titration therefore samples a one-dimensional input trajectory through the multidimensional apoptotic signaling network. The measured threshold *U*_*c*_ identifies where this trajectory crosses the network separatrix and is therefore used as a one-dimensional proxy for *u*_*c*_. Assuming that the mapping between the experimental input and the reduced model threshold is approximately proportional over the conditions examined, *u*_*c*_ ∝ *U*_*c*_, then 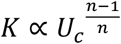. Thus, in the high *n* limit, we can take the experimentally measurable quantity *U*_*c*_ to be proportional to *K*. The dependence of wave speed on *U*_*c*_ therefore becomes:

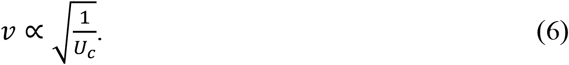

This means that if other factors (the positive feedback doubling speed and the diffusion coefficient) remain constant, the wave speed should approximately scale with the inverse square root of the activation threshold.

To test this relationship, we replotted the data from Fig. 2f as a log-log plot; according to Equation 6, the data should fall on a straight line with a slope of −0.5. As shown in Fig. 4a, the equation agreed well with the overall trend in the data and the fitted slope was −0.62 (95% CI, −0.77 to −0.47).

**Fig. 4:**
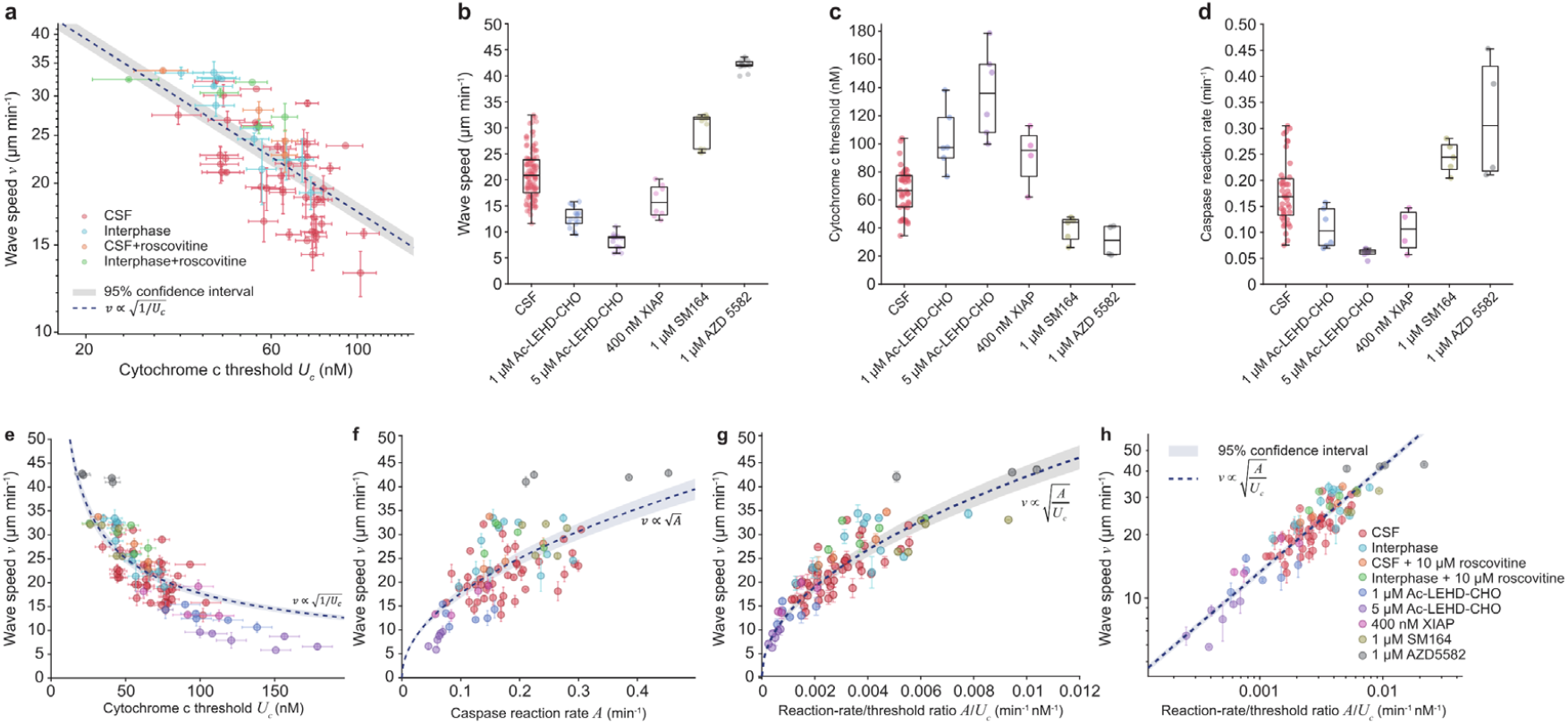
Apoptotic trigger wave speed scales with the cytochrome c activation threshold and the caspase reaction rate. a, Cytochrome c threshold (*x*-axis) versus apoptotic trigger wave speed for CSF and interphase extracts with or without the addition of the Cdk inhibitor roscovitine (10 µM). These are the same data as those shown in Fig. 2f except plotted on logarithmic axes. A fit of the equation 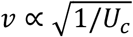 (dashed line) with the 95% confidence interval is shown. b, Boxplots of apoptotic wave speeds in CSF extracts (N = 113), or CSF extracts treated with 1 μM Ac-LEHD-CHO (N = 16), 5 μM Ac-LEHD-CHO (N = 16), 400 nM XIAP (N = 8), 1 μM SM164 (N = 13), or 1 μM AZD5582 (N = 14). c, Boxplots of the cytochrome c activation thresholds in CSF extracts (N = 42), or CSF extracts treated with 1 μM Ac-LEHD-CHO (N = 6), 5 μM Ac-LEHD-CHO (N = 6), 400 nM XIAP (N = 4), 1 μM SM164 (N = 5) and 1 μM AZD5582 (N = 4). P-value from rank-sum test is labeled on top. d, Boxplot of caspase reaction rates in CSF extract with different drug treatments, as in Fig.4b. P-value from rank-sum test is labeled on top. e, Relationship between apoptotic trigger wave speed (*v*) and the cytochrome c activation threshold (*U*_*c*_). The error bars of wave speed represent the standard deviation within each frog from 1 to 4 replicates, and the error bars of the cytochrome c represent the step size of the cytochrome c gradient used in the experiment. The experimental conditions are color coded as denoted in (h). The dashed curve is a fit of the relationship 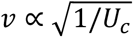 to the data. f, Relationship between apoptotic trigger wave speed (*v*) and the caspase reaction rate (*A*). The error bars of speed represent standard deviation within each frog from 1 to 4 replicates. The shaded areas represent the 95% confidence interval. The experimental conditions are color coded as in (h). The dashed curve is a fit of the relationship 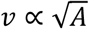 to the data. g, h, The quantity 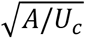 predicts the trigger wave speed. Panel g is a linear plot; panel h a log-log plot. The dashed curve (g) and dashed line (h) are fits of the relationship 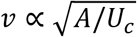 to the data. The experimental conditions are color coded as in (h).

To extend this experimental test further, we treated extracts with drugs, peptides, and proteins targeting different steps in the caspase-3 activation pathway to alter wave speed, reaction thresholds, and/or reaction speeds, and asked whether the relationship 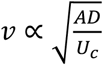 held under all these conditions. We assumed that these perturbations did not substantially change the relevant diffusion coefficient. We measured the maximum caspase-3/7 reaction rate by adding excess Z-DEVD-R110 to fully apoptotic extract and fitting the cleavage dynamics with Michaelis-Menten kinetics (Fig. S2 a-d, see methods), and we measured the caspase threshold as described for Fig. 2. Inhibiting caspase-9 with Ac-LEHD-CHO or adding XIAP reduced the trigger-wave speed and maximal caspase reaction rate and increased the cytochrome c activation threshold (Fig. 4b-d). Conversely, the XIAP antagonist SM164 increased wave speed and caspase reaction rate and lowered the activation threshold. AZD5582, a broader IAP antagonist, produced a stronger acceleration, with higher caspase reaction rate and lower *U*_*c*_ (Fig. 4b-d).

We fitted the relationships 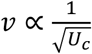 and 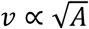 to the experimental data. Wave speed was found to decrease with *U*_*c*_ (Fig. 4e) and increase with *A* (Fig. 4f), as expected, but there were quantitative discrepancies between the data and the equations. For example, the data points with the highest thresholds *U*_*c*_ all fell below the theoretical curve in the wave speed versus *U*_*c*_ fit, and the points with the lowest thresholds all fell above it (Fig. 4e). The converse was true for the wave speed versus *A* fit (Fig. 4f). The wave speed data was better accounted for by the two parameters together, 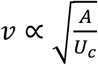 (Fig. 4g), and a log-log plot of *v* versus *A*/*U*_*c*_ yielded a linear relationship with a slope close to the theoretical value of 0.5 (slope = 0.47, 95% CI, 0.42 to 0.51) (Fig. 4h). Thus, *U*_*c*_ and *A* together explained most of the observed wave-speed variation across cell-cycle states and pharmacological perturbations, and the theoretical expression 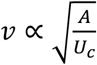 was found to hold for a range of experimental conditions.

Finally, we calculated the residuals after fitting the equation log_10_*v* = *β* + 0.5log_10_*A* − 0.5log_10_*U*_*c*_ to the wave speed data (Fig. S3a, b). Residuals of the fitted speed were similar across conditions (Fig. S3b), and pairwise comparisons after Benjamini-Hochberg false-discovery-rate correction did not identify significant residual differences among conditions (Fig. S3c). Thus, *U*_*c*_ and *A* together and the scaling relationship defined by Equation 6 explain most of the observed wave-speed variation across cell-cycle states and pharmacological perturbations.

## Discussion

Luther’s equation 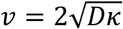, which is identical in form to the Fisher-KPP equation, provides a simple benchmark for the speed of a trigger wave in terms of measurable properties of the system that generates the trigger wave. However, when we examined the speeds of apoptotic trigger waves in CSF-arrested versus interphase *Xenopus* egg extracts, we found a difference in speeds that could not be accounted for by differences in *D* or *κ* (Fig. 1). This led us to derive an extended form of Luther’s law that acknowledges that as a trigger wave mediator diffuses into a new volume, the system’s positive feedback may not be able to initiate exponential activation until the concentration of the mediator exceeds some non-negligible threshold. Indeed, experiments indicated that in *Xenopus* extracts, a substantial threshold concentration of cytochrome c must be exceeded before any production of active caspase-3/7 can be detected (Fig. 2).

To account for a threshold in the trigger wave speed expression, we began with a one-variable reaction-diffusion equation with a Hill function in its positive feedback reaction term, and then simplified the reactions further to a piecewise-linear one-variable model (Fig. 3). For such a system the wave speed 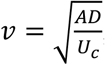, where *A* is a measure of reaction speed, *D* is the diffusion coefficient, and *U*_*c*_ is the threshold for the bistable system. This expression agrees well with numerical simulations of the original one-variable Hill equation system provided that the Hill exponent is reasonably large (Fig. 3).

We then tested whether this extended form of Luther’s law could account for the different apoptotic wave speeds in interphase versus CSF *Xenopus* extracts. The answer was yes; moreover, a number of perturbations that increased or decreased the sensitivity of the extract to an apoptotic stimulus (cytochrome c) could also be rationalized by the extended form of Luther’s law (Fig. 4).

This scaling relationship may extend beyond apoptosis to other biological trigger waves driven by switch like activation dynamics, including cell cycle waves^7^, ERK/MAPK activation waves^9,40^, and ferroptosis propagation^13^. However, the predicted inverse-square-root scaling should be expected only when the wave is driven by a thresholded reaction with high Hill coefficient rather than a pulled wave dominated by infinitesimal leading-edge growth.

## Methods

### Preparation of *Xenopus* egg extracts

Female *Xenopus laevis* (LM00531, NASCO) used in this study were at least 3 years old. Animal care and handling followed a protocol (APLAC-13307) approved by the Institutional Animal Care and Use Committee (IACUC) of Stanford University. Cytostatic factor (CSF)-arrested and interphase-arrested egg extracts were prepared using established protocols^6,29,30^. Briefly, female frogs were primed with 100 IU of pregnant mare serum gonadotropin (PMSG) (BioVendor, LLC) and ovulation was induced by injection of 500 IU of human chorionic gonadotropin (hCG) (Millipore-Sigma). Frogs were kept in egg-laying buffer (100 mM NaCl, 2 mM KCl, 1 mM MgSO_4_, 2.5 mM CaCl_2_, 0.5 mM HEPES and 0.1 mM EDTA, pH 7.4) overnight. Eggs were collected and dejellied in 2% cysteine solution (w/v) in buffer (100 mM KCl, 0.1 mM CaCl_2_, 1 mM MgC1_2_). For CSF-arrested extracts, eggs were washed with CSF extract buffer (100 mM KCl, 50 mM sucrose, 10 mM potassium HEPES pH 7.7, 5 mM EGTA pH 7.7, 2 mM MgCl_2_, and 0.1 mM CaCl_2_). For interphase extracts, eggs were washed with extract buffer (100 mM KCl, 50 mM sucrose, 0.1 mM CaCl_2_, 1 mM MgCl_2_, 10 mM HEPES, pH 7.7). The eggs were packed by centrifugation at 200 ×g for 1 min followed by 600 ×g for 30 s. Leupeptin (L2023, Sigma), pepstatin (P5318, Sigma), chymostatin (C7268, Sigma), cytochalasin B (C6762, Sigma), and nocodazole (HY13520, MCE) were added to concentrations of 10 µg ml^−1^ each. Cycloheximide (01810, Sigma) was added to interphase extracts to a concentration of 100 µg ml^−1^. Eggs were lysed and fractionated by centrifugation at 16000 ×g for 15 min at 4°C, and the middle cytoplasmic layer was collected by needle puncture.

Unless otherwise indicated, extracts were supplemented with 5 µM Z-DEVD-R110 (CPC Scientific Inc.) as a fluorogenic caspase-3/7 sensor, and 10 µM bortezomib (Selleck Chemicals) to suppress background fluorescence. Apoptotic extracts were prepared by adding horse cytochrome c (Sigma Aldrich) to 1 μM. For perturbation experiments, Ac-LEHD-CHO (CPC Scientific Inc.), SM-164 (S7089, SelleckChem), and AZD5582 (HY12600, MCE) were dissolved in DMSO and added to extracts at a final DMSO dilution of 1:1000 (v/v). XIAP (Sinobiological) was dissolved in water and added to extracts at a final dilution of 1:100 (v/v). Final reagent concentrations are indicated in the corresponding figures and figure legends.

### Apoptotic trigger wave assay and image analysis

Cytoplasmic extracts were loaded by vacuum into Teflon tubing with a 300 µm inner diameter at room temperature. To initiate apoptosis from one end, the tubing was dipped for 10 s into an apoptotic extract. The activated tubes were transferred to a FluoroDish (World Precision Instruments, FD5040-100) and covered with mineral oil. Fluorescence imaging was performed at room temperature on a Leica DMi8 inverted epifluorescence microscope equipped with a 5× objective.

An empty-field image was collected for shading correction. Background-corrected images were stitched with a custom algorithm. Image analysis was performed in MATLAB R2025b. For each time frame, fluorescence was projected perpendicular to the long axis of the tube by maximum-intensity projection. At each axial position, the wave arrival time was defined as the first time point at which fluorescence reached half of its local maximum. Wave speed was estimated by linear regression of arrival time versus position. Only fitted regions spanning at least 1 mm and having R^2^ > 0.95 were used for speed estimation.

### Cytochrome c activation threshold measurements

To determine the cytochrome c concentration required for apoptotic activation, extract aliquots containing Z-DEVD-R110 were treated with graded concentrations of cytochrome c and monitored with a fluorescence plate reader. For each extract, the fluorescence value 20 min after cytochrome c addition was normalized to the minimum and maximum response across the cytochrome c titration series. A Hill equation 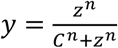 was fitted to the stimulus-response data and the parameter *C* of the fitted curve was used as the cytochrome c activation threshold, *U*_*c*_. For activation frequency plots, an extract was classified as activated at a given cytochrome c concentration if the endpoint fluorescence (at 20 min) exceeded half of the maximum response in that experiment. Uncertainty in *U*_*c*_ was taken as the cytochrome c concentration step size used in the titration.

### Protein diffusivity measurement in *Xenopus* egg extract from FCS

Protein diffusivity in *Xenopus* egg extract was measured by fluorescence correlation spectroscopy (FCS) as described previously^41^. Measurements were performed on a Zeiss LSM 980 laser scanning confocal microscope at the Stanford University Cell Sciences Imaging Facility (CSIF; RRID:SCR_017787) using a C-Apochromat 40× water-immersion objective. The pinhole was calibrated with the Zen3 software automatic pinhole-adjustment routine. Coverslip *z*-position and thickness were measured in reflectance confocal mode, and FCS measurements were collected approximately 60 µm above the coverslip surface. Atto 488-carboxylic acid in water (100 nM; diffusion coefficient *D* = 400 μm^2^/s) was used to calibrate the observation volume. Alexa-488 conjugated bovine serum albumin (100 nM) is added to *Xenopus* extracts for diffusivity measurement. Photon counts were collected for 60 s using an MBS 488/639 filter and a GaAsP PMT detector. Autocorrelation curves were analyzed with the FCS module in Zeiss Zen 3 software and fitted with a three-dimensional anomalous-diffusion model:

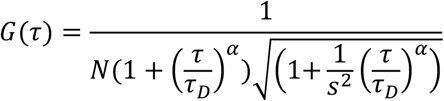

Here, *G*(*τ*) is the normalized autocorrelation curve, *N* is the particle number, *τ*_*D*_ is the characteristic diffusion time, *α* is the diffusion-mode parameter and *s* is the optics structural parameter. The effective Brownian diffusion coefficient was calculated by 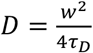 where *w* is the spot size determined from the Atto 488 calibration.

### Measurement of caspase-3/7 reaction rates in apoptotic extracts

Caspase-3/7 was fully activated by adding 500 nM cytochrome c to a *Xenopus* egg extract and incubating the extract for 30 min. A calibration curve relating fluorescence intensity to cleaved Z-DEVD-R110 concentration was generated by adding known concentrations of Z-DEVD-R110 to fully activated extract and incubating for an additional 30 min; the relationship was linear throughout the concentration range (Fig. S2a). For kinetic measurements, 20 µM Z-DEVD-R110 was added to extracts containing fully activated caspase-3/7, and fluorescence dynamics were recorded. Because the total caspase-3/7 concentration in *Xenopus* egg extracts is approximately 200 nM^36^, substantially lower than the substrate concentration, cleavage of Z-DEVD-R110 was modeled using Michaelis-Menten kinetics:

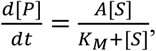

where [*P*] is the concentration of cleaved substrate and [*S*] is the concentration of uncleaved substrate, *A* is the maximum reaction rate, and *K*_*M*_ is the Michaelis constant. Given [*P*_*total*_] = [*P*] + [*S*], and assuming that when *t* = 0, [*P*] = *p*_0_, the integrated solution is:

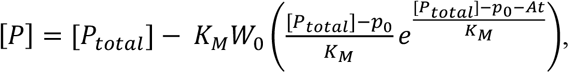

where W_0_ is the principal branch of the Lambert W (Product Log) function. Fluorescence traces were fitted in MATLAB R2025b using the Curve Fitting Toolbox with a nonlinear least squares method and trust-region algorithm. All fitted parameters were constrained to be non-negative. Because direct nonlinear estimation of *A* produced large standard errors, the reaction-rate parameter used in subsequent analyses was estimated from the initial linear regime of the kinetic data. Calibrated fluorescence traces were converted to product concentration, and the available substrate concentration was calculated as total substrate concentration estimated from nonlinear fitting minus the instantaneous product concentration. The instantaneous cleavage rate, 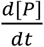, was calculated numerically. The saturated region with available substrate < 0.4 µM and the nonlinear region with available substrate > 8 µM were excluded. The slope of the remaining initial linear regime was used as the experimental estimate of the caspase-3/7 reaction rate parameter *A*.

### Obtaining the caspase-3/7 doubling time from Luther’s equation

The apparent caspase-3/7 doubling speed, *κ*, was estimated using a previously described approach^20^. We let *k*_*R*_ denote the reaction-rate constant for Z-DEVD-R110 cleavage by active caspase-3/7 and *k*_*BG*_ denote the background cleavage rate of the Z-DEVD-R110 sensor. The normalized fluorescence *I*(*t*) from caspase-3/7 catalysis can be expressed as:

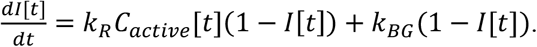

Assuming the caspase-3/7 activity follows a logistic temporal profile:

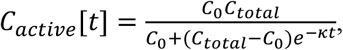

And solution of the fluorescence equation gives:

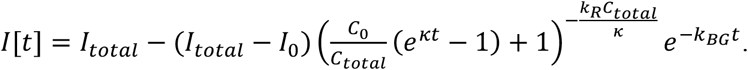

Here *I*_0_ and *I*_*total*_ are the initial and total normalized Z-DEVD-R110 fluorescence values, *C*_0_ is the initial active caspase-3/7 concentration, and *C*_*total*_ is the maximum active caspase-3/7 concentration. The reaction rate constant *k*_*R*_ was computed from *C*_*total*_ and the maximum caspase-3/7 reaction rate *A* as 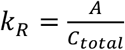. For each experiment, fluorescence traces from individual positions along the tube were normalized, aligned to their first half-maximum crossing time, averaged, and fitted to the model above. Fitting was performed in MATLAB R2025b using the Curve Fitting Toolbox with a nonlinear least squares method and the trust-region algorithm. The fitting range for *κ* was 10^−3^ to 5 min^−1^, *k*_*BG*_ was 0 to 0.05 min^−1^, *C*_0_ was 10^−5^ to 10 nM, *C*_*total*_ was 100 to 400 nM, *I*_*total*_ was 0.9 to 1.1, and *I*_0_ was −0.05 to 0.05. Fitted spatial regions were at least 1 mm long and had R^2^ > 0.95.

### Obtaining parameters for the Hill-type reaction term from time course experiments

The temporal fluorescence trace at each position was normalized and aligned to the first half-maximum crossing time. The average aligned trace, *I*[*t*], was used to estimate the normalized caspase-3/7 activity as 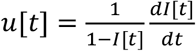 for the portion of the trace with 0.2 < *I*[*t*] < 0.8.

The activity trace *u*[*t*] was then normalized, and the phase-plane relationship between 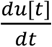 and *u*[*t*] was fitted. Regions with negative *u*[*t*] or 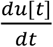were excluded.

To account for small normalization errors in the experimental activity traces, we also fitted a model with an explicit normalization parameter, *u*_*total*_:

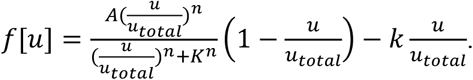

Fitting was performed in MATLAB 2025b using the curve fitting toolbox with a nonlinear least squares method. The range of the fitted parameters was 0 to 2 for *A*, 0 to 1 for *K*, 1 to 12 for *n*, 10^−6^ to 0.1 for *k*, and 0.5 to 2 for *u*_*total*_. The fitted values and 95% CI of the fitting were *n* = 5.2 (95% CI 4.4 to 6.2), *K* = 0.18 (95% CI 0.15 to 0.21), *k* = 1.1*e* − 5 (95% CI 1*e* − 6 to 3.0*e* − 3), *A* = 0.065 (95% CI 0.050 to 0.079), and *u*_*total*_ = 1.0053 (95% CI 0.90 to 1.17). Because *k* contributed little during the rapid activation phase, its confidence interval was broad. With *u*_*total*_ fixed at 1, we also performed fitting with 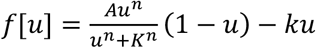, and the fitted values and 95% CI of the fitting were *n* = 5.2 (95% CI 4.4 to 6.2), *K* = 0.18 (95% CI 0.17 to 0.19), *k* = 1*e* − 6 (95% CI 1*e* − 6 to 2.7*e* − 3), and *A* = 0.065 (95% CI 0.062 to 0.069).

### Steady states for the Hill-type reaction equation

The reaction term for the one variable reaction-diffusion system included a Hill-type activation function with weak inactivation:

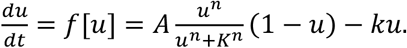

The variable *u* represents active apoptotic mediator; for example, *u* could represent active caspase-3, which is derived from a finite pool of inactive procaspase-3. We normalized the concentrations so that active plus inactive caspase/procaspase-3 equals 1. The rationale for the inclusion of a Hill function in the activation rate expression is that Hill functions are often reasonable approximations to complex biological responses. The parameter *A* is the activation rate constant, *K* is the *EC*50 of the Hill term (and is related to the threshold concentration of *u* required to trigger apoptosis), *n* is the effective Hill exponent, and *k* is the (small) inactivation rate constant. This is identical to the reaction system considered by Schlomann et al.^39^ except that we included the factor (1 − *u*) in the activation rate term.

We can obtain the steady states of the system by setting *f*[*u*] = 0 and solving for *u*. For appropriate parameter choices, there are three steady states. The first steady state, which we call *u*_*low*_, is found when *u = 0*. This represents the low caspase-activity living state, and this steady state is stable. A stable steady state with high caspase activity*—*the apoptotic state*—*is also present. If *K* is relatively small, so that the Hill function is approximately equal to one at this apoptotic steady state, then the value of *u* at this steady state is given by 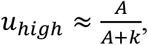 which simplifies to 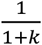 after rescaling time by A. The third steady state is unstable and lies between the other two. If this steady state is close to *u* = 0, we have (1 − *u*) ≈ 1, and:

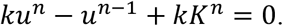

Since both *k* and *u* are small, we have *u*^*n*−1^ = *kK*^*n*^, which gives

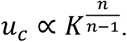

The same relationship was derived by Schlomann and colleagues for their somewhat different system^39^.

### Estimates of trigger wave speed from a piecewise-linear reaction-diffusion equation

The full reaction-diffusion equation treats the local caspase-3/7 activity as a normalized variable *u*(*x, t*), ranging from an inactive low state to an activated high state. The experimentally inferred reaction term was highly switch-like, consistent with a large effective Hill coefficient (*n* ≈ 5; Fig. 3c-e). We therefore started from the following Hill-type reaction-diffusion equation:

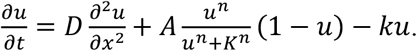

Here *D* is the effective diffusion coefficient of the propagating apoptotic mediator, *A* is the activation rate constant for the positive feedback reaction, *K* is concentration of *u* at which the Hill function is half-maximal, *n* is the effective Hill exponent, and *k* is a weak effective inactivation or loss rate. The dimensions are:

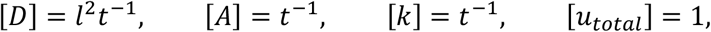

where *l* represents the length unit and *t* is the time unit.

If the Hill coefficient is sufficiently large, the activation term can be approximated by a Heaviside step function centered on *u* = *K*:

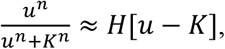

This gives a piecewise-linear sharp-threshold model:

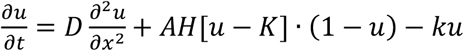

This equation suggests that ahead of the front, where *u* is below *K*, the positive feedback is not activated, and the activity only diffuses and decays. Behind the front, where *u* is above *K*, the positive feedback is on and drives the system toward the activated state. This results in a stable, propagating wavefront.

We can use the activation rate 1/*A* to set the time scale and the diffusion-activation length 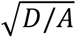 to set the length scale, and define three new variables:

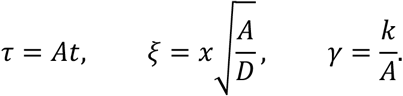

Substituting these expressions into the dimensional PDE gives

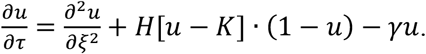

The dimensional speed *v* and dimensionless speed *c* are therefore related by:

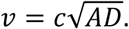

In the inactivate region ahead of the wavefront, where the Heaviside function evaluates to zero, the non-dimensionalized PDE reduces to:

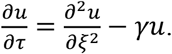

The inactive steady state of the reaction term corresponds to *u*_*inact*_ = 0.

In the activated region, the non-dimensionalized PDE becomes:

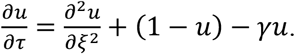

The activated steady state (*u*_*act*_) is obtained by setting the reaction term in this equation to zero.

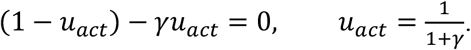

We now seek a traveling wave solution moving to the right at constant speed *c*. This means that position and time are related, and we can define a new variable *U* such that:

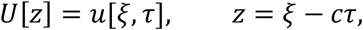

The boundary conditions are:

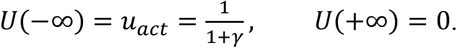

The front location can be defined by the threshold-crossing condition:

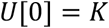

Here, the activated region is *z* < 0, the inactive region is *z* > 0.

Substituting the traveling-wave ansatz into the non-dimensionalized PDE gives:

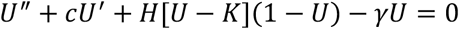

For the region ahead of the wavefront, the local activity is below threshold. This equation becomes:

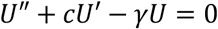

Trying an exponential solution *U* = *e*^*az*^ yields the characteristic equation: *a*^2^ + *ca* − *γ* = 0. The two roots are:

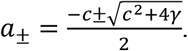

The solution ahead of the front must decay to zero as *z* goes to positive infinity. Therefore, the admissible root is the negative root, denoted by 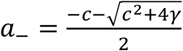, and the solution ahead of the wavefront satisfying *U*(0) = *K* is *U*_*a*_(*z*) = *Ke*^*a*−*z*^.

The derivative at the threshold point is:

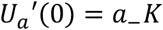

The characteristic equation also gives the relationship between *a* and *c*:

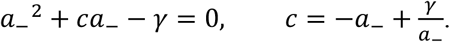

For the region behind the wavefront, the local activity is above threshold. The non-dimensional ODE becomes:

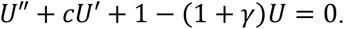

Subtracting the activated plateau by writing W = *U* − *u*_act_ gives:

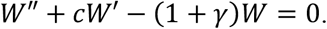

The characteristic equation is: *b*^2^ + *cb* − (1 + *γ*) = 0, and the two roots are:

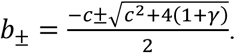

A perturbation from the activated state must decay as *z* goes to negative infinity. Therefore the admissible root is the positive root, denoted by 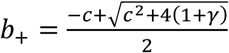:

The behind-front solution satisfying the conditions *U*[−∞] = *u*_act_ and *U*[0] = *K* is therefore *U*_*b*_[*z*] = *u*_act_ − *Qe*^*b*+*z*^, where *Q* = *u*_act_ − *K*. The derivative at the threshold point is:

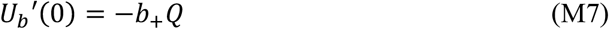

The characteristic equation also gives relationships between *b* and *c*:

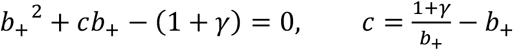

Since the derivative of *U* must match at the threshold, we have:

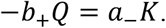

Substituting *b*_+_ = −*a*_−_*K*/*Q* and *c* = −*a*_−_ + *γ*/*a*_−_ into *b*_+_^2^ + *cb*_+_ − (1 + *γ*) = 0, yields:

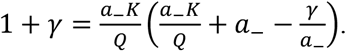

After collecting terms and using:

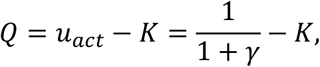

the decay rate ahead of the front is

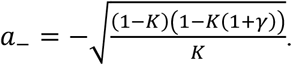

Since *c* = −*a*_−_ + *γ*/*a*_−_, the dimensionless trigger-wave speed is:

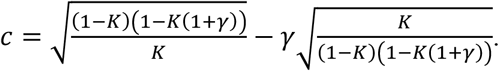

Recall that 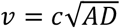 and 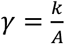. Therefore the dimensional wave speed is:

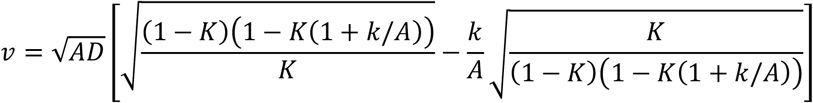

In the experimentally relevant limit where inactivation is weak compared with activation, 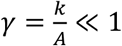, and the speed reduces to:

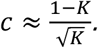

If *K* is small compared with the activated state, then 1 − *K* ≈ 1, and the leading scaling becomes 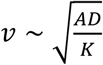.

From previous method section we have:

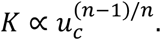

Here *u*_*c*_ denotes the unstable steady state of the one-variable reaction model. The experimentally-determined cytochrome c activation threshold *U*_*c*_ is used as a proxy for this value. For large values of the Hill coefficient *n*, (*n* − 1)/*n* ≈ 1, and so the model threshold *K* is approximately proportional to the experimentally measured activation threshold:

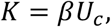

where *β* is the calibration factor converting the measured cytochrome c threshold into the normalized model threshold. Substituting this into the speed expression we have:

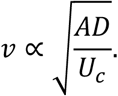

This relationship was experimentally tested in Fig. 4a,g,h.

### Numerical simulations

Numerical simulations of wave speed were performed in MATLAB R2025b using the PDE Toolbox. The threshold parameter *K* was scanned either linearly from 0.02 to 0.5 in increments of 0.02 or logarithmically from 2^−6.5^ to 2^−1^ in increments of 2^0.25^. The Hill coefficient *n* was set to 2, 4, or 6, or fixed at 5, as indicated. The inactivation parameter *k* was set to 0.001, 0.01, or 0.1, or fixed at 0.001. Unless otherwise stated, *D* = 1 and *A* = 1. Simulations were performed in one spatial dimension with reflective no-flux boundary conditions at both ends. Initial conditions were set to the high steady state in a seed region at the left boundary and to the low steady state elsewhere. The integration used relative tolerance 1e-5, absolute tolerance 1e-7. For each position, the wave arrival time was determined by linearly interpolating the time at which *u* crossed the midpoint between the low and high stable states. Local speeds were calculated from the distance and crossing-time differences between adjacent positions, and the reported wave speed was taken after the local speed stabilized.

## Supplemental Figures and Legends

**Fig. S1:**
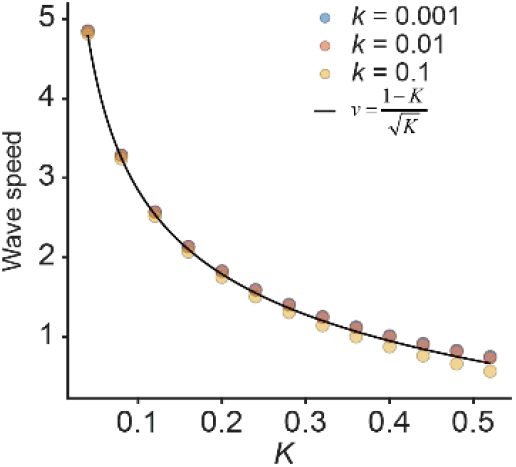
Contribution of degradation rate to wave speed. Simulated wave speeds as a function of activation threshold (*K*) with varying degradation rates (*k*) are shown as dots. Theoretically predicted wave speed base on the piecewise-linear model (which yields 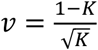) is shown as a black line (*D* = 1, *A* = 1, *n* = 5).

**Fig. S2:**
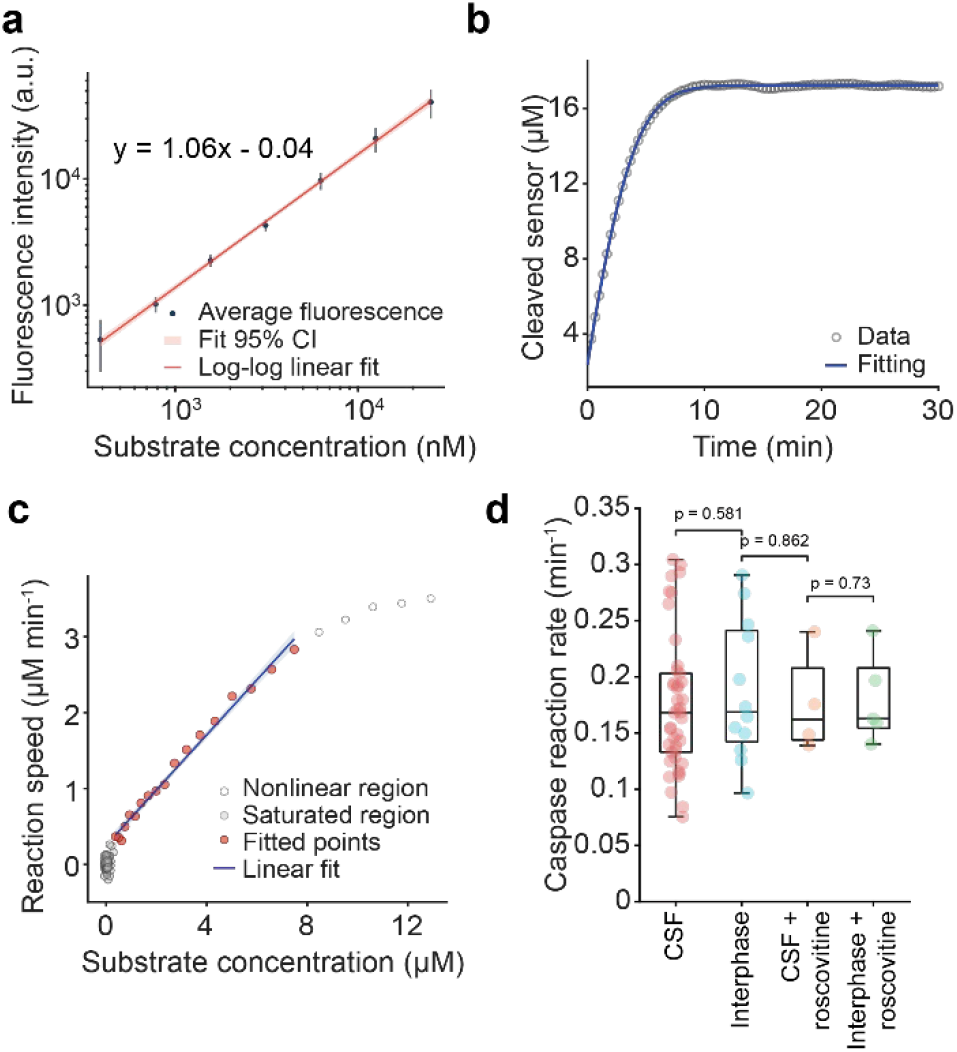
Measuring the maximum reaction rate in fully apoptotic extracts. **a**, Calibration curve of the fluorescence plate reader. Fluorescence intensities in extracts with different Z-DEVD-R110 concentrations were measured after a sufficiently long time following apoptosis induction, showing a near linear relationship between fluorescence intensity and the amount of cleaved sensors. **b**, A representative fluorescence intensity time course after apoptosis activation (black dots) and the fitted curve obtained from integrating Michaelis-Menten kinetics (blue lines, methods). **c**, Relationship between instantaneous reaction rate and available substrate concentration from experimental data (blue dots) and linear fitting (red line), highlighting that the reaction rate is linearly related to substrate concentration. **d**, Caspase reaction rates in CSF extracts (red, N = 42 frogs), interphase extracts (blue, N = 12 frogs), CSF extracts treated with 10 μM roscovitine (orange, N = 4 frogs), and interphase extracts treated with 10 μM roscovitine (green, N = 5 frogs).

**Fig. S3:**
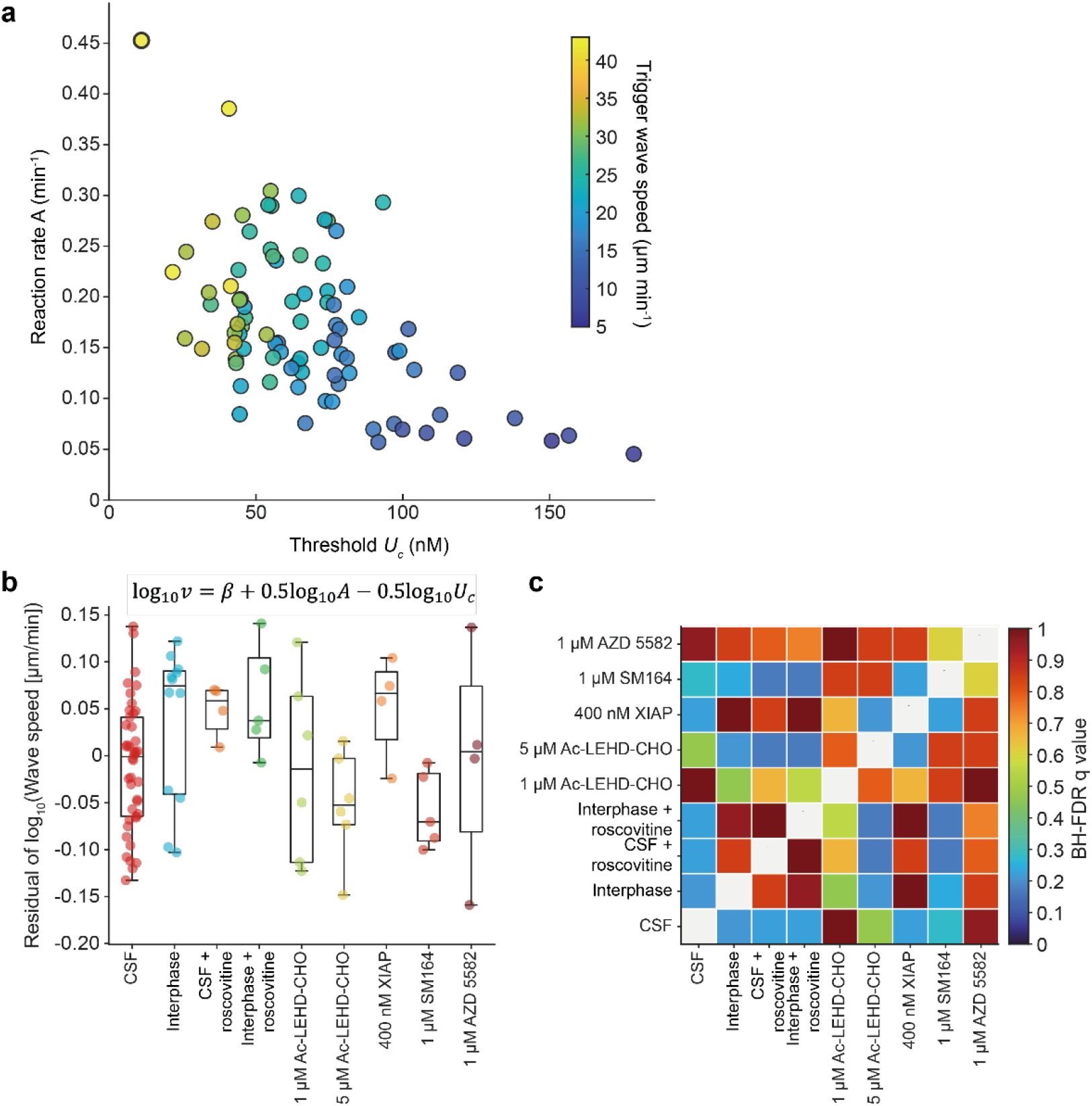
Estimating trigger wave speed from activation threshold and caspase reaction rate. **a**, Relationship between cytochrome c activation threshold *U*_*c*_, caspase reaction rate *A* and wave speed *v*. The data points correspond to experimental measurements with CSF or interphase-arrested extracts treated with various inhibitors as below. The colors denote the corresponding trigger wave speeds for the measurements. Note that the wave speed increases as the threshold decreases (right to left) and as the reaction rate increases (bottom to top). **b**, Distribution of speed residuals. An overall value for β in the equation shown was calculated from the experimental data, and the residuals were calculated for each experimental wave speed determination. R^2^ = 0.82. Data are from experiments in CSF extracts (N = 42 frogs), interphase extracts (N = 12 frogs), CSF extracts with roscovitine (10 µM) (N = 4 frogs), interphase extracts with roscovitine (10 µM) (N = 5 frogs), CSF extracts with: 1 μM Ac-LEHD-CHO (N = 6 frogs), 5 μM Ac-LEHD-CHO (N = 6 frogs), 400 nM XIAP (N = 4 frogs), 1 μM SM164 (N = 5 frogs) and 1 μM AZD5582 (N = 4 frogs). **c**, Benjamini-Hochberg false discovery rate (BH-FDR) for pairwise comparisons of speed residuals among conditions. No pairwise comparison was significant after BH-FDR correction.

